# Cell-to-cell Mathematical Modeling of Arrhythmia Phenomena in the Heart

**DOI:** 10.1101/2020.07.28.225755

**Authors:** Gabriel López Garza, A. Nicolás Mata, G. Román Alonso, J. F. Godínez Fernández, M. A. Castro García

**Affiliations:** Mathematics Department, Universidad Autónoma Metropolitana Iztapalapa Ciudad de México, México; Electric Engineering Department, Universidad Autónoma Metropolitana Iztapalapa, Ciudad de México, México

**Keywords:** Fibrillation, Fluttering, Arrhythmia, Pseudo-Electrogram, Mathematical modeling

## Abstract

With an aperiodic, self-similar distribution of two-dimensional arrangement of atrial cells, it is possible to simulate such phenomena as Fibrillation, Fluttering, and a sequence of Fibrillation-Fluttering. The topology of a network of cells may facilitate the initiation and development of arrhythmias such as Fluttering and Fibrillation. Using a GPU parallel architecture, two basic cell topologies were considered in this simulation, an aperiodic, fractal distribution of connections among 462 cells, and a chessboard-like geometry of 60×60 and 600×600 cells. With a complex set of initial conditions, it is possible to produce tissue behavior that may be identified with arrhythmias. Finally, we found several sets of initial conditions that show how a mesh of cells may exhibit Fibrillation that evolves into Fluttering.

## 1. Introduction

For the sake of mathematical simplicity, we define only two types of arrhythmia in excitable media. One type is known as *Fluttering* and is related to reentrant waves of excitation, which remain in a self-perpetuating steady state. The second and more complex type of arrhythmia considered in this article is known as *Fibrillation*. Meanwhile, Fluttering is adequately described employing continuous or cell-to-cell modeling; the Fibrillation phenomenon is more difficult to simulate with deterministic models. Since the first research papers, some authors, [5], [38], considered that Fibrillation could only be approached mathematically on a statistical basis, mainly due to the random distribution of anastomosis fibers in the heart. This standpoint is still in use [33], [34], [36], [42], but the primary mechanisms of flutter and Fibrillation are not fully understood. The researchers still have incomplete knowledge of how arrhythmias, such as ventricular Fibrillation, begin and develop. In opposition to the statistical basis thesis, we present a deterministic model in which we introduce complexity in the cellular network geometry as a factor for the generation of arrhythmias. In a network with simple topology, we produce Fibrillation by adding a set of complex initial conditions in a completely deterministic set of ordinary differential equations.

In this article, we study some of the consequences obtained by modeling weakly connected networks through different distributions of excitable cells within the mesh, what we call *the geometry of the network*. In this context, we argue in subsection 3 why cell-to-cell modeling fits better than the continuous model, at least to model the arrhythmia. We will illustrate in the Methods section 2, that neither the diffusivity provided by partial differential equations nor by the cell-to-cell coupling requires a complex dynamics in the cells to produce fibrillation and flutter phenomena. Elliptic-type operators give diffusivity in continuous mathematical modeling and also in cell-to-cell modeling using *weakly coupled variables* (see section 3.0.1). Nevertheless, we show *in silico* that fibrillation and Fluttering can be modeled even by using the simplest excitable cell models including only a few variables and “realistic models of heart cells” and compare the silico experiments of both realistic vs. few variables models [22], [24].

The main difference between flutter and Fibrillation, according to the classic definitions [38], is the randomness of Fibrillation as opposed to the regularity of flutter. Randomness precludes sharp, well-defined wavefronts. One contribution of our work is to introduce some degree of complexity (the tiling of Figure 5) instead of randomness to present an *in silico* phenomena, which can be identified with Fibrillation. We simulated Fibrillation in a simple Chessboard geometry in a mesh of 60 × 60 and a mesh of 600 × 600 cells by introducing a complex set of initial conditions. This Fibrillation is achieved with both models, two variables and Nygren model of the human heart. Another novelty in the present paper is that, contrary to the commonly established, Fluttering can be produced at a cellular level by a dynamic obstacle formed with a few cells and also by fixed non-dynamical obstacles (for a definition see section 2.2.1). Finally, we found several sets of initial conditions that show how, even in the simplest mesh of cells, they may exhibit Fibrillation that evolves into Fluttering. This phenomenon that is well known in medical literature, for the best of our knowledge, is for the first time shown with human heart cell models.

## 2. Methods

### 2.1. Individual Cell models

In this work, we use two-variable models of excitable cells [1], [4], [12], [24], as well as a physiologically accurate model of Nygren and coworkers [27]. The idea of using two variables vs. many variables models is to extract the properties of a net of cells that depend only on the excitable media and do not depend on the limitations of individual cell models.

The models of excitable cells included here, as usual, go through four stages [40]: resting, exciting, excited, and refractory states; also, the models of coupled cells provide solitary waves flexible enough to flutter and fibrillate. In this way, the models represent observables in real tissue to some extent (for a mathematical definition of *observable* see section 3.0.1). The convenience and relevance of utilizing more complex models of individual cells is discussed in sections 5 and 6.

#### 2.1.1. Realistic Models

There are many physiologically accurate models of excitable cells in the Heart, among them: Courtemanche et al., [8], Nygren et al., [27]; Lindblad et al., [21]. In this paper, we use Nygren et al. model (N), taking into account that the N model reconstructs action potential data that represent recordings from human atrial cells. In the N model, the sustained outward K^+^ current determines the duration of the action potential (AP). On the other hand, the AP shape during the peak and plateau phases is determined by transient outward K^+^ current, I_*sus*_, and L-type Ca^2^ current. The N model has 29 variables: 12 transmembrane currents, a two-compartment sarcoplasmic reticulum (SR), and restricted subsarcolemmal space for calcium dynamics handling and calcium buffering.

Regarding the number of variables, simulating a system of 360 000 cells, as we do in this work, is somehow onerous in computational terms. For example, 360 000 times 29 gives a system of ODE of 104 400 variables. For instance, simulating 15 seconds with step-size of the order of milliseconds may illustrate what we mean by “onerous computationally”. To accomplish this task, we employed parallel architecture using Nvidia GPU (RTX 2080 Ti), and we developed it with C-CUDA libraries. The Procedures and Algorithms that we used are described in Nicolas and coworkers [25]. To solve the ODE systems, we implemented a Runge-Kutta numerical method of order four with absolute error tolerances of 10^−6^.

#### 2.1.2. Modeling Atrial fibrilation

For the Nygren model, we used the data in Cherry et al. [7] and references therein to simulate electrophysiological changes that occur as a result of sustained Atrial Fibrillation. Specifically, I_*Ca,L*_ is decreased 30 percent of its original value, and I_*to*_ and I_*Kur*_ are both decreased to 50 percent of their original values. As mentioned in Cherry et al., APs are triangular in morphology at all cycle lengths and are shorter under these conditions. Additionally, rate adaptation is largely abolished. The reduction in rate adaptation shown by the model is in agreement with some experimental studies of chronic Atrial Fibrillation (AF) and tissues obtained from right atrial appendage tissue of patients with chronic AF (see references therein [7]). Low conductivity of cells in the heart is associated with ischemia [18], and in experiments, conductivity may be lowered pharmacologically by heptanol [5].

Finally, to simulate Fibrillation as those in Figure 15, we found sets of initial conditions by implementing a random search in both models N and B. In generating in the net the initial conditions, we alternate stimulated cells with refractory state cells. These complex sets of initial conditions in continuous media modeled with partial differential equations represent a discontinuous function and, therefore, highly improbable sets. However, modeling cell-to-cell such complex sets of initial conditions is not improbable since its discontinuity is inherent to the heart tissue structure. Discontinuity in the real tissue is another argument that favors ODE modeling over PDE modeling. Summarizing, we consider two types of initial conditions: (a) One small connected set of exciting cells surrounded by refractory cells; (b) Many small islands of exciting cells scattered throughout the entire net mixed with refractory cells. The interested reader may obtain our data on the sets of initial conditions under request to the corresponding author.

A vast difference exists among two-variable models and realistic ones, regarding, for instance, the number of observable phenomena. Nevertheless, all ordinary differential equations models we used have four states. A rest state corresponding to a minimum value of the AP variable; an exciting state, which corresponds to a negative derivative of the AP profile; an excited state, corresponding to the maximum of the AP profile; and a refractory state, associated to a positive derivative of the AP profile. In figures 1 and 2(b), the rest state is represented in the deepest blue color, and the excited state is represented in the darkest red.

**Figure (1).**
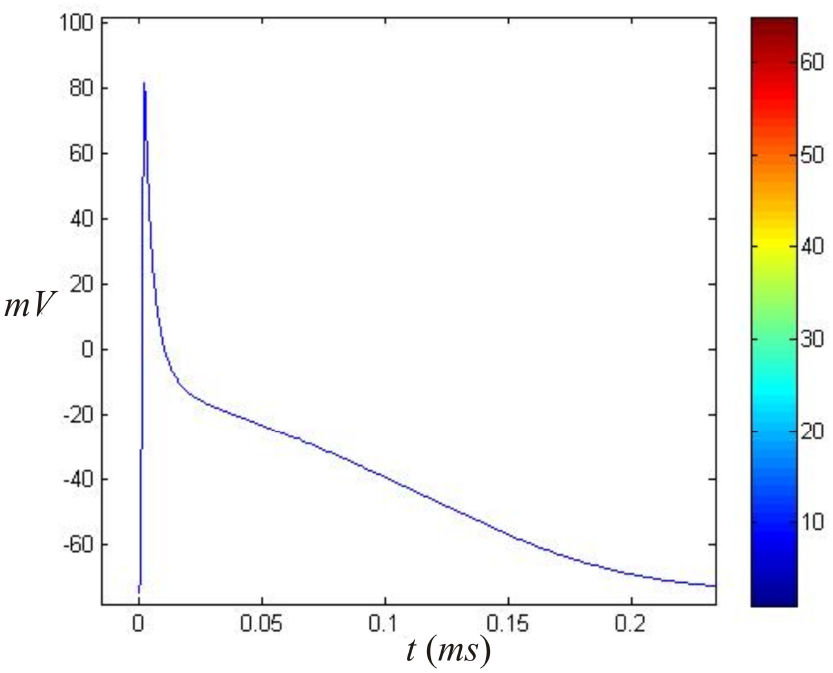
Color code of the AP corresponding to Nygren’s model. In the video captures and videos, the deep blue color corresponds to a rest state and deep red to a maximum of the action potential in the Nygren et al. cell model.

**Figure (2).**
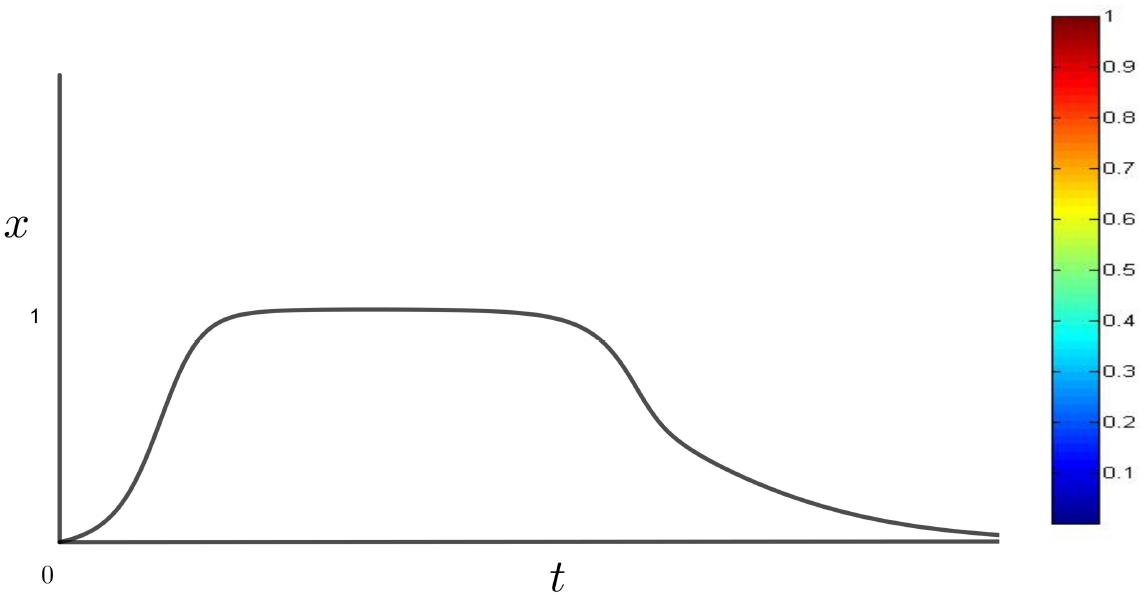
The AP corresponding to the Barkley model of equation (1), recall that *x* and *t* are adimensional for this model. Each value of *x* of the plot in the left corresponds a color to each height. The color bar corresponds to the Barkley model’s states in the videos and two-dimensional plots shown in this paper. Deep blue corresponds to rest state *x*(*t*) =0 and the darkest red corresponds to *x*(*t*) = 1.

#### 2.1.3. Simple ordinary differential equations models

For this part, although only the experiments with the Barkley [4] model are reported, the Fitzhugh-Nagumo model and the Aliev-Panfilov model whose description is elsewhere [12], [24], [1] were also subject of experimentation. Since results obtained for Fluttering and Fibrillation are similar to those obtained with the Barkley model, Fitzhugh-Nagumo, and Aliev-Panfilov, we do not include plots of the last two, and from now on we will only refer to the Barkley model when we talk about two-variable models. If a suitable geometry of the cell system is introduced (see section 2.2), it is possible to represent Fibrillation and flutter phenomena with all these models. They are two-variable models, as is well known, and they are dynamical bi-stable systems. For these systems, the existence of limit cycles is well established in the mathematical theory, and even *analytical* approximations of physiologically relevant limit cycles in a region between heteroclinic trajectories are possible to calculate [17].

The Barkley model used in this paper is the following

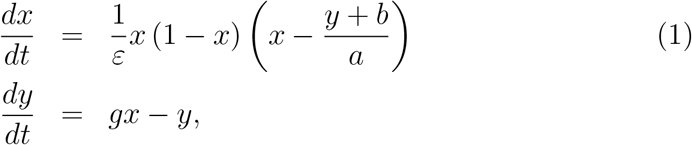

where *a, b, g, ε* are fixed parameters. Figure 2 shows an AP of Barkley model with initial conditions *x*(0) = 0.4, *y*(0) = 0. The variable *x* in this article corresponds to an adimensional voltage, and may be identified with the variable *V* of the Nygren model.

### 2.2. Cell-to-cell Nets Geometry

The geometry in cellular systems can be determined by considering the geometry of the individual cells and how they are connected. For example, the working cells in the auricula in the heart are mostly cylindrical and are connected in a way that favors the longitudinal transmission of Action Potentials [35]. By comparison, brain cells have extremely branched forms, and their connections can reach a complexity that is far from being understood in its entirety [31]. In the following system,

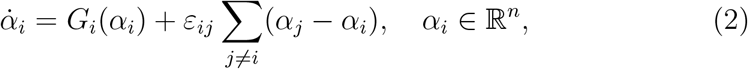

where *n* represents the number of variables of each cell. The geometry of the network (and hence the diffusivity) is determined by the values of *ε_ij_* different from zero. So, equation (2) can be written as a vectorial equation with *α* = (*α*_1_, …, *α_n_*), *G*(*α*) = (*G*_1_(*α*), …, *G_n_*(*α*)). So, as an illustration, for two variables with the Barkley model in equation (1), the system (2) has the form

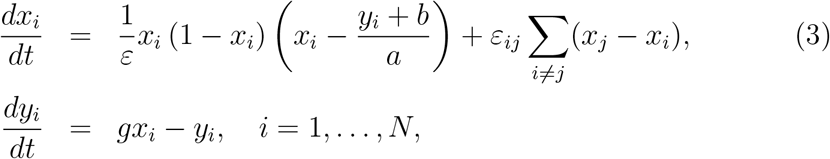

where *N* is the number of cells in the system and *ε_ij_* = 0 for unconnected cells. Notice that only the *x_i_* variables are coupled to each other as corresponds to variables related to the Action Potential in the heart’s cells. Similarly, for the N model of atrial cells of 28 variables, only the voltage is coupled.

In practice, the use of rectangular and cubic matrices are the most commonly used [9], [13], [28], [41], without considering the complex geometry of the cytoarchitecture of the network that exists in the tissues of living beings. Systems made of systems of equations of the form (2) are known as *weakly connected networks* (WCN) and have a vast number of applications in neurophysiology [14]; cardiology [41], [9], [28]; and many other sciences. In general, WCN have applications in every system of cells connected through gap junctions, such as those in atrial tissue in the human heart.

#### 2.2.1. Obstacles

There are two types of obstacles to be considered: dynamical and static. Dynamical obstacles are formed by individuals or groups of cells in a refractory or excitable or excited state. Static obstacles are formed by objects that do not change in time. They may correspond to fibroblast, adiposity in real tissue, or dead tissue due to heart attacks. In this article, we consider only dynamic obstacles.

#### 2.2.2. Tiling

One of the central thesis of this work is that, under certain circumstances, an intricate connection between cells is essential in the generation of flutter and Fibrillation. We take as a paradigm of cell connections, hence the intrinsic geometry of the cells, those of the working cells in the auricula, and the ventricle in the heart. In the literature, histological studies of heart’s cells are available [35]. Nevertheless, mathematical models, including the real geometry of the cells, are more scarce. Spach and Heidlage [36, Fig. 1] give a schematic representation myocardial architecture of 33 cells in a two-dimensional array. Following their representation, Figure 4 depicts a two dimensional model of cells, but our model is not based in real cells as in [36] but in a distribution generated by an aperiodic tiling called “ Table” which we describe below. After the cells’ connectivity is fixed, it is possible to establish a correspondent cell geometry, as in Figure 4. Note that the random-like distribution of the cells is not for real; in Figure 4 are represented in the up and left corner in Figure 5. This distribution can be verified, noting that the code of colors corresponds to the same cells, meaning green for cells with six connections, pink for cells with five connections, and so on. Observe that in our figure, cell connections occur only in the vertical edges where most of the standard electrical coupling between cells have place [35].

The values *ε_i,j_* different from zero in equation (2) give an adjacency matrix between cells which represents the geometry of the entire net. In this paper, we use a “Table” distribution of connections among cells. A *Table* is a polygon belonging to the class that can be tiled by a finite number of smaller, congruent copies of itself (see [32], where the properties of the “Table” as tiling-dynamical-system are studied). We used this tiling for the following reasons. a) It is an aperiodic tiling of the plane so that some degree of complexity is intrinsic in the adjacency matrix. b) The tiling is self-similar, so it does possess a fractal structure. c) Each cell is connected to an average of 4.86 cells, which is a good 2D approximation compared with an average of 9.1 reported from experimental data measures by Hoyt et al. [15] for three-dimensional structures. Besides, the connectivity approaches that of the cells in the Spach diagrams of Figure 1 in [36], which represents a sample of two-dimensional tissue cells with average two-dimensional connectivity of 4.66 cells. d) The “Table” aperiodic setting provides the more simple arrangement in the authors’ opinion, which satisfies the mentioned properties.

Figure 5 shows a tiling (see Figure 5) used in our in silico experiments. This arrangement is only one sample of an infinite number of such aperiodic tilings. It is necessary to assign each cell a number to set a matrix corresponding to the tiling in Figure 5. Then, once the assignment is completed, the adjacency matrix can be settled down. Algorithms to construct the *Table* and other aperiodic tilings are well known, but finding algorithms to set the associated adjacency matrix of such tilings is still an open problem, to the best of the authors’ knowledge. The recursive procedure for the self-similar aperiodic tiling is shown in Figure 3.

**Figure (3).**
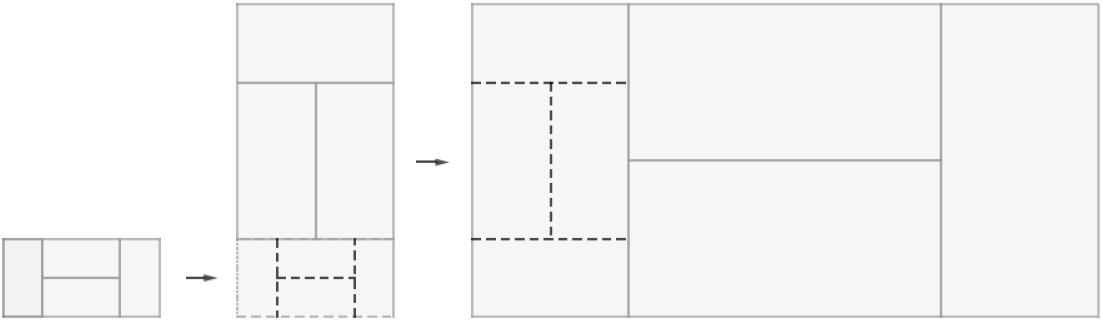
The procedure presented in the Figure is repeated several times to produce the tiling of Figure 5. Note the fractal structure obtained.

### 2.3. Flutter, Fibrillation, and pseudo-EG mathematical definitions

In this paper, we depart from the classical definitions of Flutter and Fibrillation in use to formulate the following definition that applies to the rest of the article.

#### Definition.

Flutter and Fibrillation are reentrant waves of excitation which remain in a self-perpetuating steady-state; flutter having a periodic (or nearly periodic) pseudo-electrogram (pseudo-EG), and Fibrillation having a non-periodic pseudo-electrogram. Alternatively, we call Fibrillation a steady-state, self-perpetuating pattern of systems of cells without a defined front or back wave.

#### 2.3.1. Mathematical Pseudo-EG

Following the classical definitions, we consider flutter and Fibrillation as self-generating phenomena. Fluter is considered a periodic wave of waves contrary to Fibrillation, which is considered a highly complex non-periodic wave or waves. A precise difference between flutter and Fibrillation is provided by the pseudo-EG which is calculated by the following formula:

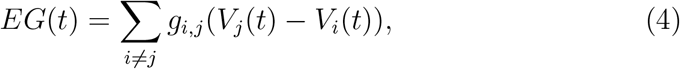

which is a non-weighted, two-dimensional version of the formula of Kazbanov et al. in [16]. Observe that the distance between cells may be taken into account handling specific weights; however, in this work this parameter is neglected since the size of the modeled tissue is small and, more importantly, the geometry and thus the real distance between cells is not modeled in our study, so that weight due to the distance may not be considered.

#### 2.3.2. Electrogram analysis

One of the most widely used techniques for analyzing fluctuations in the cardiac cycle period is the detrended fluctuation analysis (DFA). This technique has the advantage that it prevents the detection of inexistent long term correlations produced by the non-stationarity of the time series. Given the complexity of the electrograms analyzed, in this work, we used the DFA technique to determine if they present either a random behavior or a temporal structure with long term correlations, which gave us information about the propagation of the electrical signal obtained from the in silico experiments. The scaling exponents *α* obtained from the DFA analyses were evaluated as previously described by Peng et al. [29]. In short, a scaling exponent *α* near or equal to 0.5 indicates a random or uncorrelated behavior, whereas a value near or equal to 1 indicates long-term correlations in the time series; that is, current data are statistically correlated with previous data, which reflects a non-random behavior.

## 3. Theory

The concept of observable has been used through the article; next, we provide a mathematical definition. We also discuss a comparison between PDE and cell-to-cell models, which is relevant for the understanding of our results.

### 3.0.1. Observables

In mathematical modeling, the dynamics of the cells that form living beings (or that represent other excitable means) are represented by dynamic systems of form,

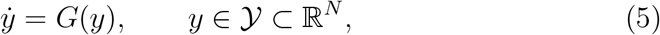

where the dot denotes the time’s derivative. So, cells are thought of as not wholly known (so far) dynamical systems, let us say

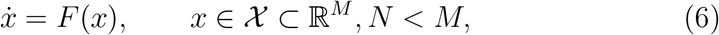

of which (5) is a representation, and scientists expect that the model *G* in some sense approaches *F*, which remains partially unknown. More formally, (5) is a model of (6) if there exists a continuous function (called observation [14]) 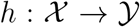 such that if *x*(*t*) is a solution of (6), then *y*(*t*) = *h*(*x*(*t*)) is a solution of (5). In practice, many information of system (6) is unknown, for instance, the dimension of the space (i. e., in this instance, the actual number of variables of the system). In many cases, a model could be a rough representation of the real system. As mentioned by Hoppenstead and Izhikevich [14], for example when (6) has a periodic solution and the model (5) is one dimensional, the observation *h*(*x*(*t*)) cannot be a solution of (5) unless *h* maps the limit cycle to a point. The existence of the function *h* and its properties are purely theoretic but allow us to speak about the relations of the real system (6) with the model in a mathematical fashion. As an example the variable *y*(*t*) = *h*(*x*(*t*)) is called an *observable*. In this article, the dimension of 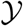 is bigger or equal than 2 so, we call *observable* to each of the *y*’s component functions.

As science advances, mathematical models of cells include an increasing number of observables and each observable with an increasing refinement following experimental data. In this way, we obtain systems of complex differential equations that include an increasing number of equations. A typical example of this phenomena is the development in the study of the sinoatrial node cells in the heart (SAN) [6], [26], (for a detailed review of the SAN mathematical models see [19]).

However, this is just the first step on the way to modeling the actual cell tissue. A second step consists of forming a system of systems of equations by coupling variables among different systems, let say *n* different systems like the following

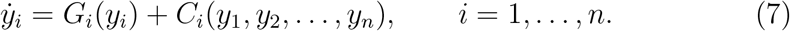

A very used example of a coupling functions *C_i_* are linear functions of the form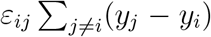 where *ε_ij_* are small (experimentally obtained) parameters and the values assigned to *j* depend on the geometry of the net. There are at least two ways of modeling excitable media. One is by utilizing partial differential equations (PDE) to represent the diffusive nature of the media. Another is by establishing a system of cells, each cell, in turn, is a system of ordinary differential equations (ODE). In ordinary differential equations, diffusion of the excitatory wave is modeled by coupling appropriate variables, for instance, the Action Potential (AP) in excitable biological cells. In this paper, we call *continuous* mathematical modeling to the first form (PDE), and the last form (ODE) is what we call *cell-to-cell modeling*.

### 3.0.2. Continuous vs. cell-to-cell modeling

Continuous mathematical modeling of anisotropic media such as ventricular tissue, normally includes fiber patterns and the continuous rotation of the fiber axis [11], so that the equations have the form:

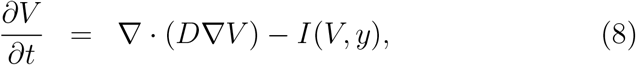

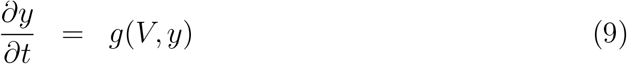

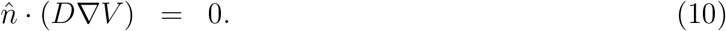

Where *V* = *V*(*t*, *x*_1_, *x*_2_, *x*_3_) is the membrane potential, (*x*_1_, *x*_2_, *x*_3_) in 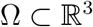, *I* is the total current through the membrane, *y* is a vector of gate variables describing the dynamics of the various currents that constitute *I*, ∇*V* denotes the gradient operator, and *D* is a conductivity tensor divided cell surface to volume ratio times the membrane capacitance of the cell. We will show that a system of equations (3) is equivalent to the system (8), (9). Note that equation (10) represents Neumann boundary conditions where 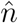 is the normal to ∂Ω. To begin with, observe that the tensor *D* is of the form

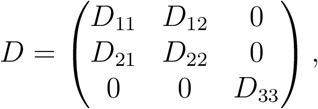

where *D_ij_* are functions of diffusivities parallel and perpendicular to the fiber, and *θ*(*x*_3_), the angle between the fiber to the axis of each plane. In the setting of [11], is easily shown that for a two-dimensional model, since *θ*(*x*_3_) ≡ 0, then *D* becomes

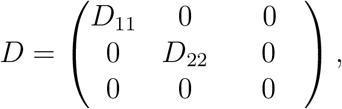

where now *D*_11_, *D*_22_ are constants, so that the elliptic operator ∇ · (*D*∇*V*) in equation (8) becomes simply

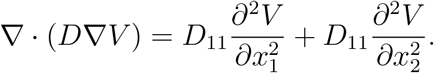

After discretization of the second partial derivatives we obtain

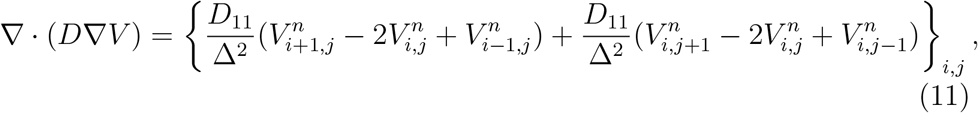

a formula which is valid for interior cells in the grid. Since in many articles including [11] the grid spacing is about Δ ≈ 200 – 300 μm which is bigger than the length of the cell, ≈ 80 *μ*m [35], is worth to mention that PDE continuous approach is, in this case, not better than cell-to-cell modeling whatsoever. Moreover, note that equation (11) corresponds to a rectangular grid of square cells (of much bigger dimensions than actual heart cells), meaning a chessboard-like geometry of the cells. In this way, a complex geometry, such as that depicted in Figure 4, cannot be represented by the elliptic operator in equation (8) since such an arrangement can not at all be represented by tridiagonal matrices such as those in equation (11).

**Figure (4).**
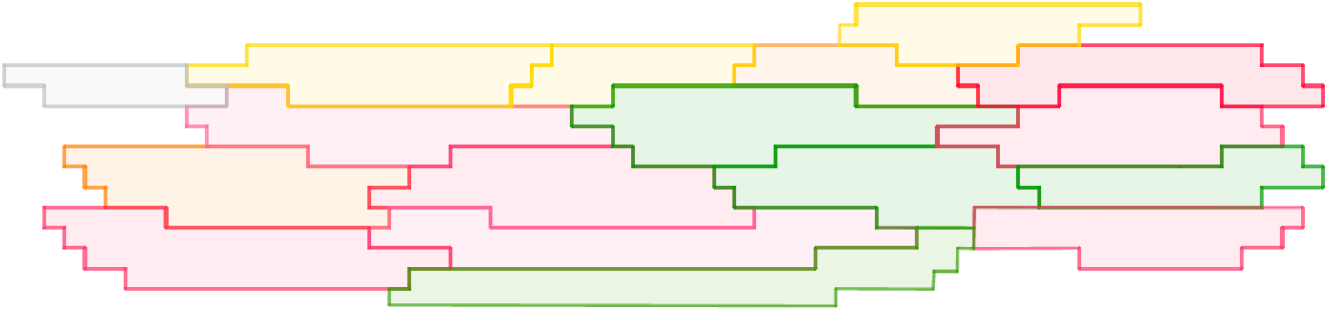
Here is shown one of many possible representations of working cells in the heart. They are not distributed randomly but following the distribution described in Figure 5, where the reader can find the color’s code. The arrangement here presented corresponds to the up and left corner of Figure 5.

A further consideration regarding the mathematics in this article must be considered. Flutter and arrhythmia will appear as solutions to systems of ODEs for a particular set of initial conditions. However, somehow the solutions appear in some fashion unpredictable since they occur after global bifurcations of the parameters given by the conductivity and of the distribution of conductivity. Hence, only after integrating the systems will emerge more of the most striking patterns of the next section in an unexpected form. Although the spirals formed by the Barkley model [2], [3], [4], and other models [10] have been extensively studied, the study of the combination of spirals, collisions of spirals, and spirals emerging after a massive dynamic blocking is still of interest regarding Fibrillation.

## 4. Results

In this paper, the waves mentioned in the definitions above are traveling waves with a defined front and back given by the ordinary differential equations of the different bistable ODE systems. As mentioned in section 2, systems with no apparent wave’s fronts and backs are noticeable. Nevertheless, a periodic pseudo-EG may appear after time in some of the systems formed with a small number of cells. While flutter is produced by the collision of traveling waves with dynamical or static obstacles at a macroscopic level as in the classic definition, there is a difference in this document with the standard definitions since microscopic (cell-to-cell) collisions are considered, and very intricate patterns may happen as, for instance, these in Figures 13 and 14.

In both types of models, two-variables and realistic Nygren model, we found that by using the tiling distribution described in Figure 5 the propagation of voltage is allowed in a very efficient way under normal conditions. Paradoxically under certain circumstances, the same topology facilitates the generation of Fluttering.

**Figure (5).**
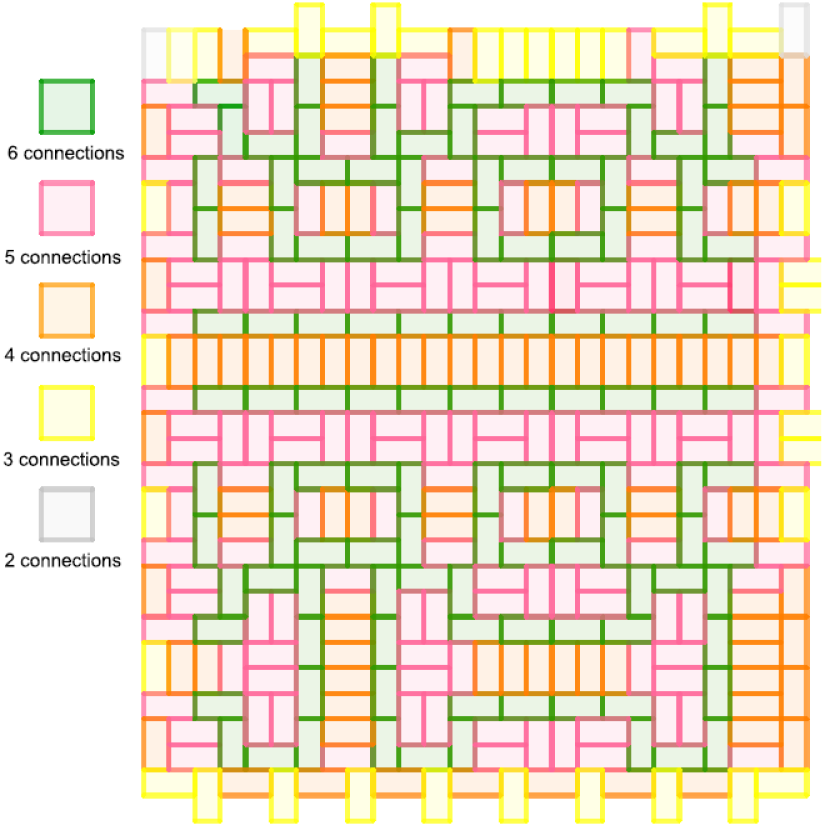
Example of a “Table” tiling showing the number of connections of each cell. Cells in green are connected with six cells, cells in pink are connected with five cells, and so on, as indicated in the figure’s left-hand side.

Moreover, in a chessboard arrange of 600 × 600 cells, we found that the generation of some Fibrillation generated by a randomly stimulated number of cells evolves to a stable multi-spiral which resembles Fluttering at least in the generation of pseudo-EG with some periodic resemblance as, for instance, in Figure 7.

We recall that we presented the description of the 29-variable model N in section 2.1.1 and a two variable model B in section 2.1.3. Given that only the Nygren model corresponds to real atrial cells in the heart, we used the DFA technique only with this model. In Table 1 we present the *α* values obtained for each in silico experiment. From subsection 2.1.2, recall that there are two types of initial conditions: (a) One small connected set of exciting cells surrounded by refractory cells; (b) Many islands of exciting cells scattered throughout the entire net mixed with refractory cells. We recall that a scaling exponent *α* near or equal to 0.5 indicates a random or uncorrelated behavior, whereas a value near or equal to 1 indicates long term correlations in the time series.

**Table (1).**
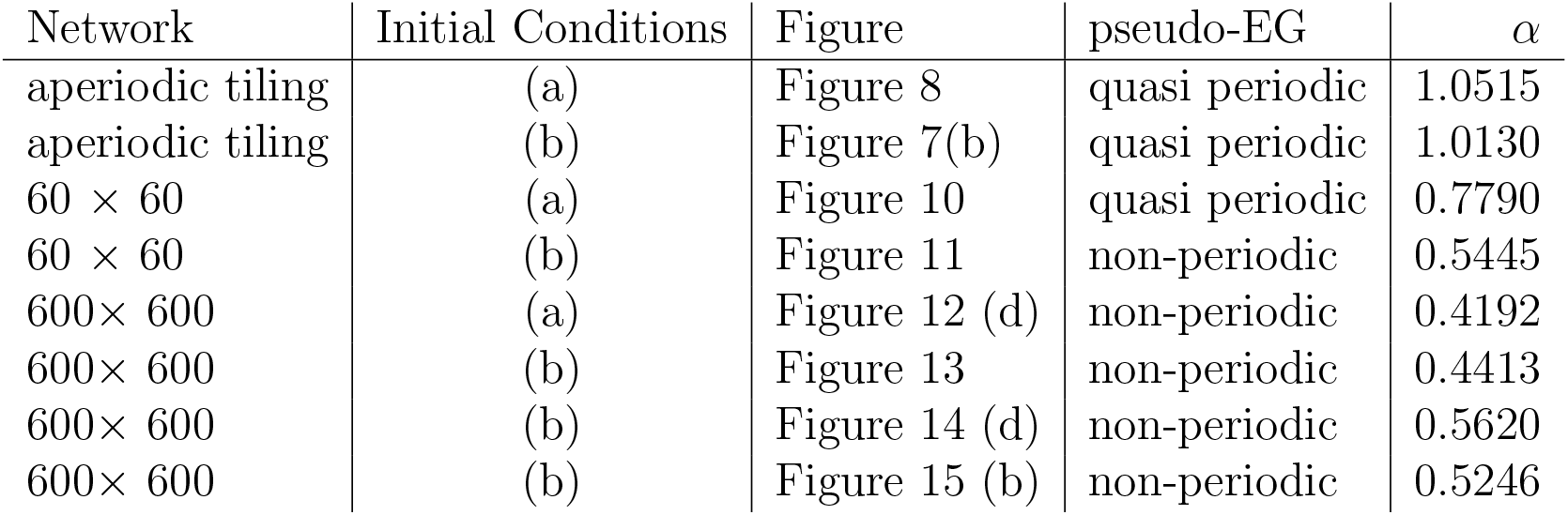
Detrended Fluctuation Analysis for N model (atrial cells)

Given the coincidences between non-realistic vs. realistic models, we conclude that the generation of Fluttering and fibrillations does depend strongly on the nature of the diffusive media, more than in the variables involved in the modeling. Nevertheless, realistic models facilitate the reduction of parameter values according to experimental data that occur during arrhythmias.

### 4.1. Fibrillation at a microscopic level case Nygren Model

In Figure 6 after a massive blocking with dynamical obstacles (initial conditions type (a)) in the aperiodic distribution of cells of Figure 5, an apparent Fibrillation becomes a Fluttering, i. e., a self-perpetuating spiral. This event is well known in the literature, but as far as we know, it is for the first time reproduced in silico.

**Figure (6).**
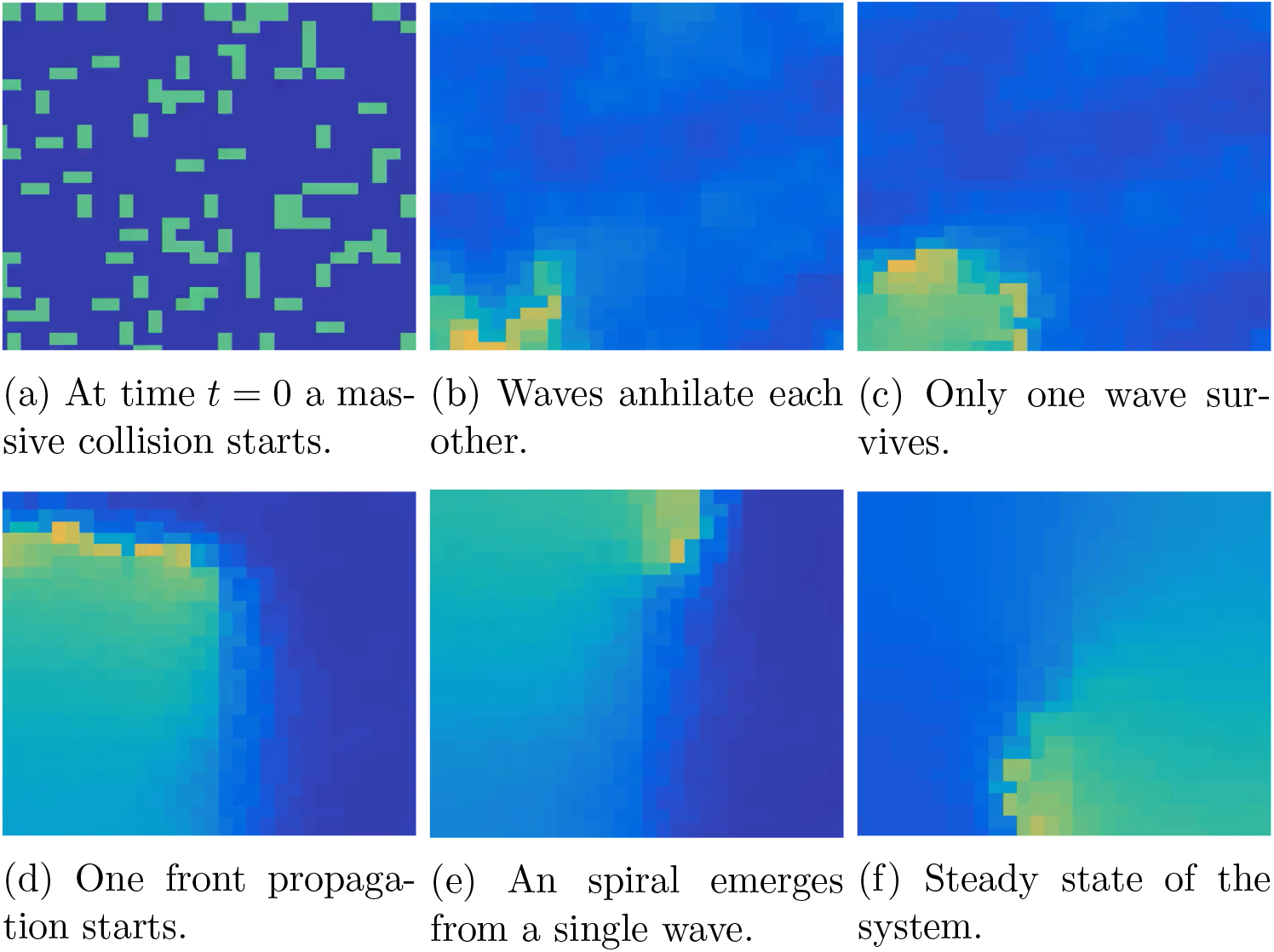
Massive blocking produces Fibrillation and evolves to Fluttering here in the tiling of Figure 5. The link to the video of the complete sequence is http://pacifico.izt.uam.mx/aurelio/.

This coincidence in the formation of spirals of the two models, one caricature (Barkley) and the other realistic (Nygren), provides us with evidence that the Fibrillation that becomes Fluttering occurs naturally in any diffusive media under appropriate initial conditions. Notice that the pseudo-EG after a non-periodic behavior from *t* = 0 to *t* < 2500 (recall that *t* is adimensional for Barkley model) resembles a periodic plot. See Figure 7.

**Figure (7).**
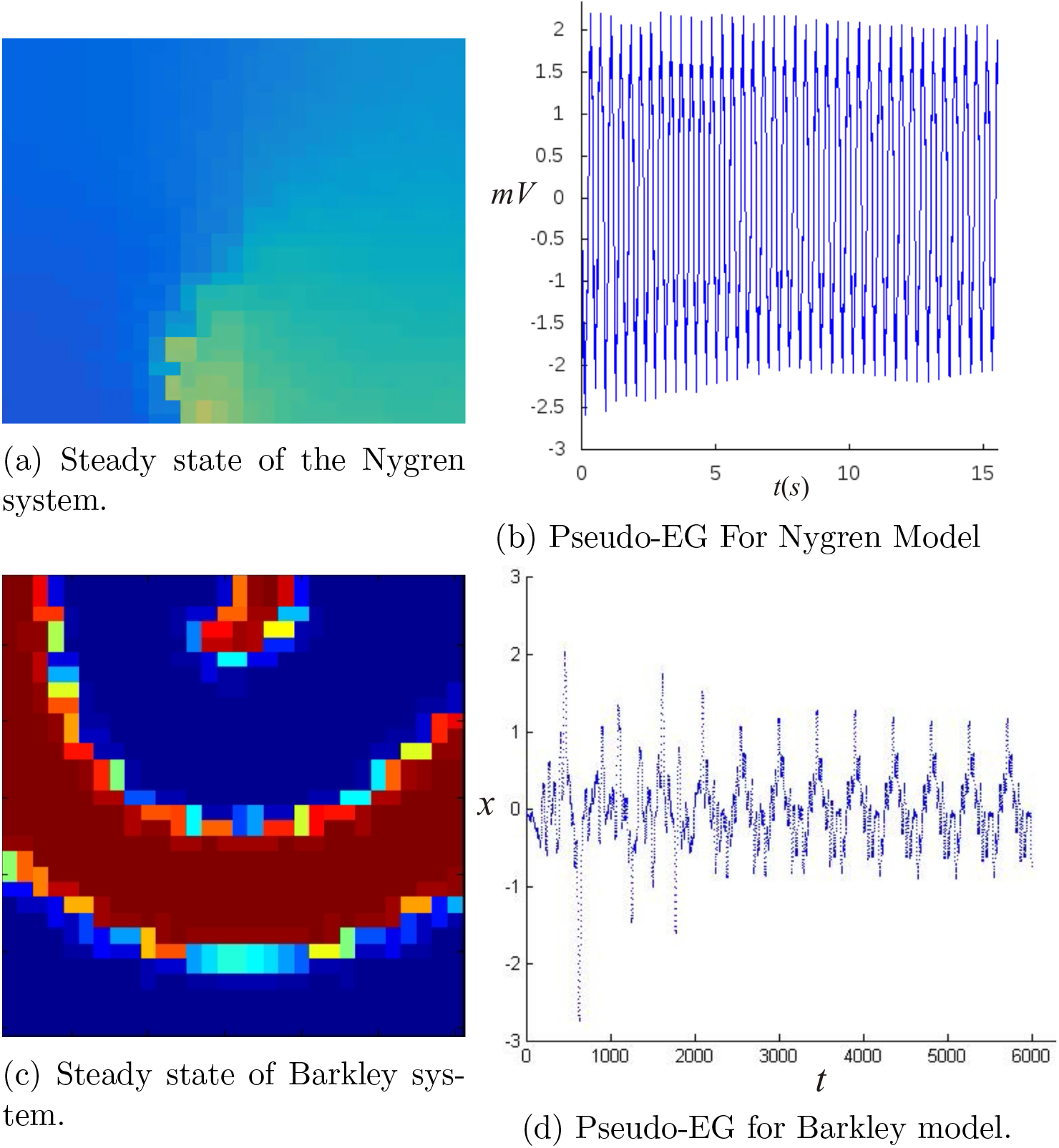
In both Nygren and Barkley models, massive blocking produces Fibrillation, which becomes Fluttering. Noticeable differences do appear in the wavelength as a consequence of intrinsic dynamics in each model, but in both of the models under certain sets of initial conditions, a periodic self-perpetuating spiral emerges. The pseudo-EG in both Figure 7(b) and 7(d) show certain periodicity.

Another interesting kind of Fluttering is produced by stimulating a small number of connected cells and blocking them with neighboring cells in a refractory state. Here, the array of cells facilitates the propagation due to its fractal nature paradoxically when a dynamic obstacle (a group of cells out of phase) prevents the wave from propagating. See Figure 8.

**Figure (8).**
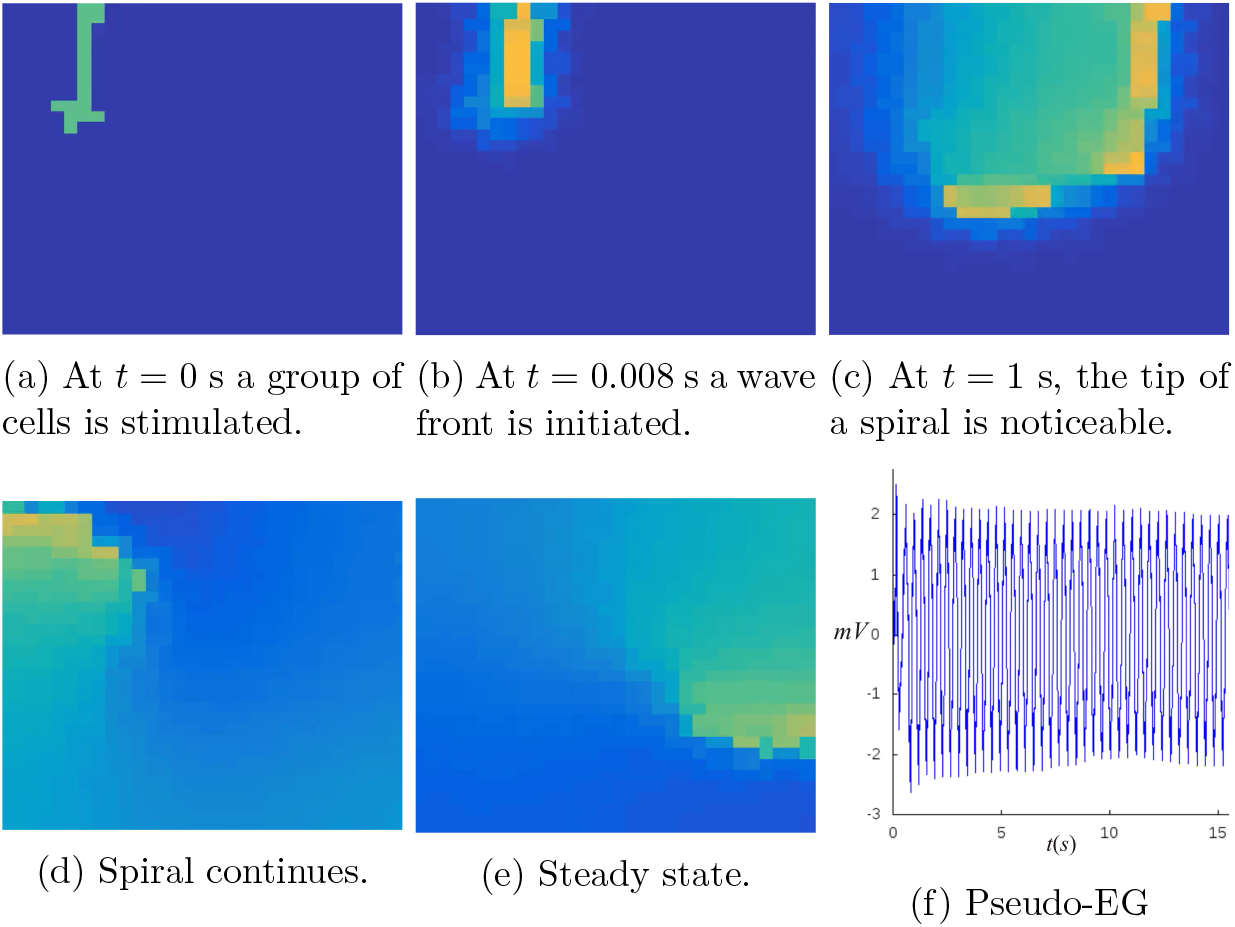
(a) A group of cells surrounded by cells in a refractory state is stimulated. (b), (c), (d), (e) A self-generating spiral which may be identified with Fluttering is apparent. (f) A quasiperiodic pseudo-EG is produced after a small, turbulent interval of time similar to that in Figure 7d.

A similar phenomenon occurs with the Barkley model, but in this case, by blocking only one cell. A single excited cell surrounded by refractory cells produces (when the network has critical connectivity) an intricate pattern of diffusion. In this way, some spirals arise due to variable cellular connectivity and low conductivity. It is worth mentioning that spirals broke into scrolls, which, according to the electrogram obtained, may be identified with Fibrillation of the system (see Figure 9).

**Figure (9).**
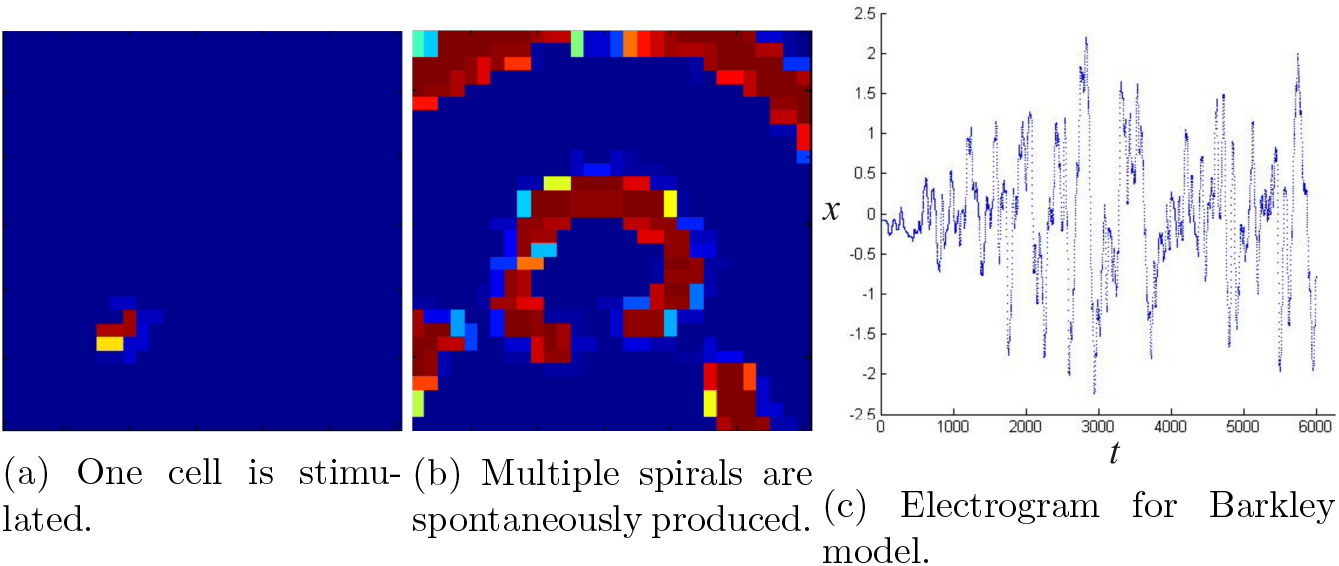
Barkley model produces several self-generating spirals by blocking one cell, with neighboring cells in a refractory state. The electrogram produces a noise-like signal which may be identified with Fibrillation.

After studying the Table array, we proceed to study a more significant number of cells but scattered in simpler arrangements. For both the Nygren and Barkley models, we started with a square of 60 by 60 cells. Furthermore, we continue with a 600 by 600 with the Nygren model. We present two types of in silico experiments. One type we block a large number of cells and scattered throughout the network, another group is stimulated. The second type, only a few connected cells, is stimulated and blocked by neighbors. We obtained the same phenomenon produced with the array. Hence a self-generating spiral is produced, see Figure 10.

**Figure (10).**
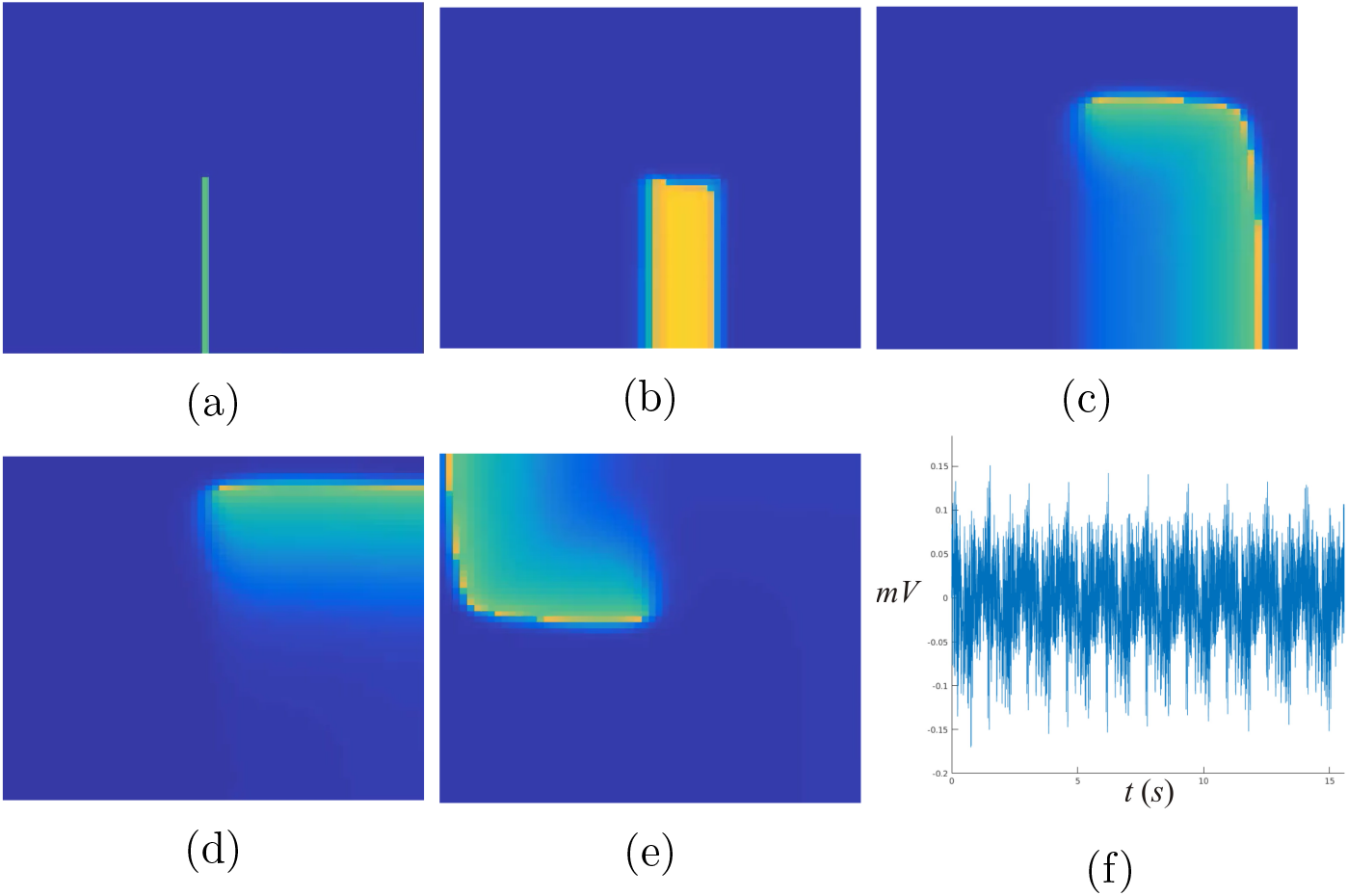
Nygren model in a 60 × 60 net of cells produces a self-generating spiral by dynamic blocking. a) At *t* = 0, a group of cells is stimulated. b) A wavefront is produced. c) A self-generating spiral appears. d) Spiral collides with the border. e) Steady-state. f) Quasiperiodic pseudo-EG.

In Figure 11 the same phenomenon illustrated in Figure 9 is represented in a 60 × 60 array of cells. A set of initial conditions alternating refractory state cells with stimulated cells produces a pattern that may be identified with Fibrillation due to the complex pattern of the pseudo-EG.

**Figure (11).**
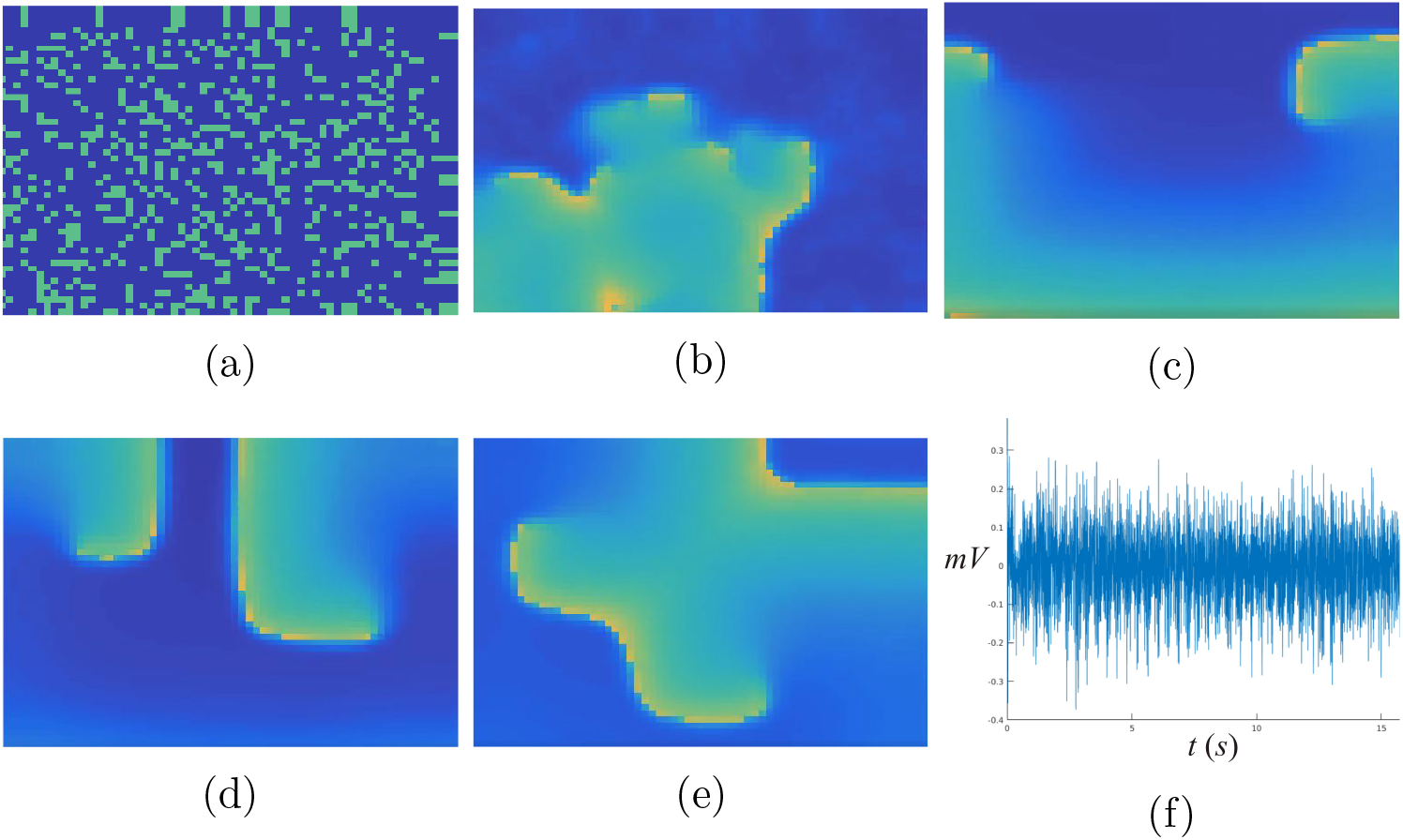
a)At *t* = 0 a massive group of scattered cells through the entire 60 × 60 net is stimulated randomly. b) Waves propagate at *t* = 0.001s. c) A couple of fronts emerge. d) Spiral after colliding produces a complex pattern e) Steady-state. f) A couple of self-generating spirals leads to a complex pseudo-EG, which may be identified with Fibrillation.

With a 600 × 600 cell array, a spiral is produced with a small group of stimulated cells surrounded by cells in a refractory state. See Figure 12. Observe that assuming that a cell has 100 *μ* m of length, a square of 360 000 cells is equivalent to 36 mm ^2^ of tissue.

**Figure (12).**
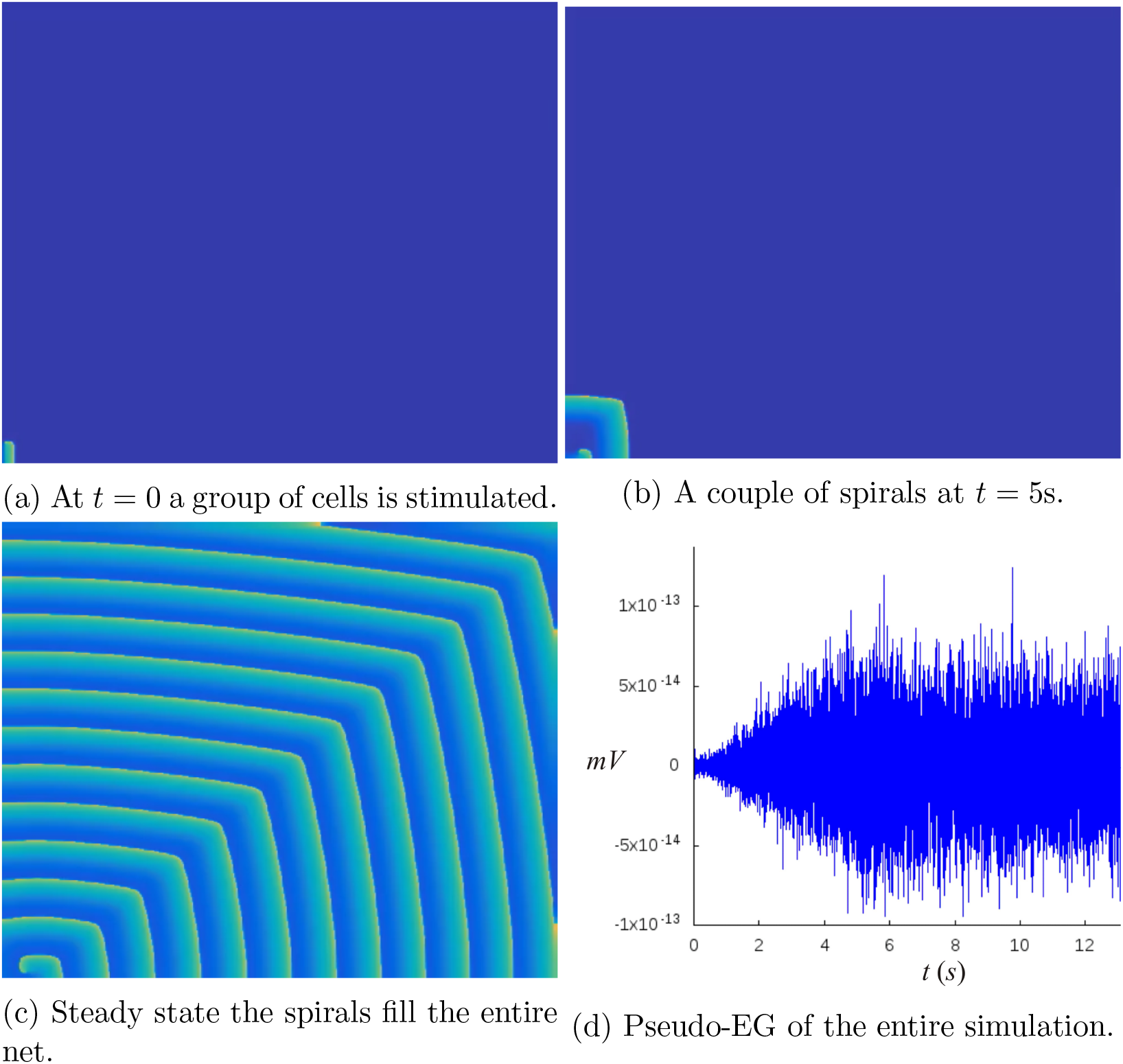
Nygren model in a 600 × 600 net of cells produces a self-generating spiral by dynamic blocking. (a) Notice the small group of stimulated cells in the left bottom corner. (b) A series of spirals emerge. (c) Steady-state. (d) Pseudo-EG of the entire simulation.

A fascinating phenomenon occurs in a 600 × 600 array when a large group of stimulated, scattered cells through the entire net is surrounded by refractory cells. Fibrillation occurs during several seconds, and under certain initial conditions, it becomes Fluttering, which persists in a stationary and complex spiral. See Figure 13.

**Figure (13).**
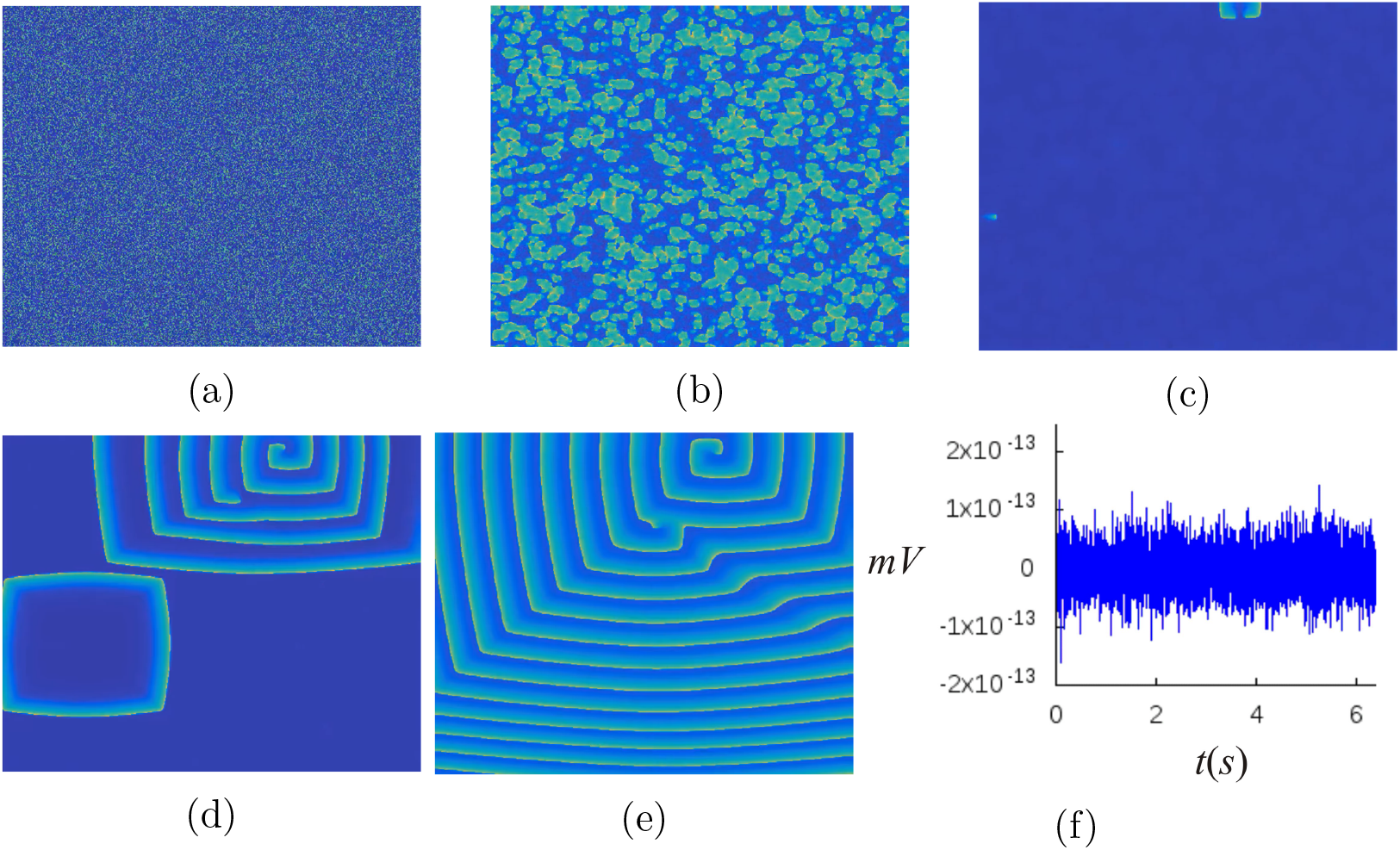
a) At *t* = 0 a massive group of scattered cells through the entire net is stimulated, but each cell is surrounded by refractory cells. b) Waves propagate at *t* = 0.08s. c) After the collisions, only two fronts survive in this particular setting. d) Spirals are formed with the remanent waves. e) A complex pattern emerges of self-generating spirals. f) Pseudo-EG of the entire simulation.

Moreover, in the 600 × 600 array, we can produce with a set of initial conditions a more complex pattern than in Figure 13 f). Many self-generating spirals resembling a micro-reentry associated with Fibrillation emerges after the collision presented with a massive blocking. See Figure 14.

**Figure (14).**
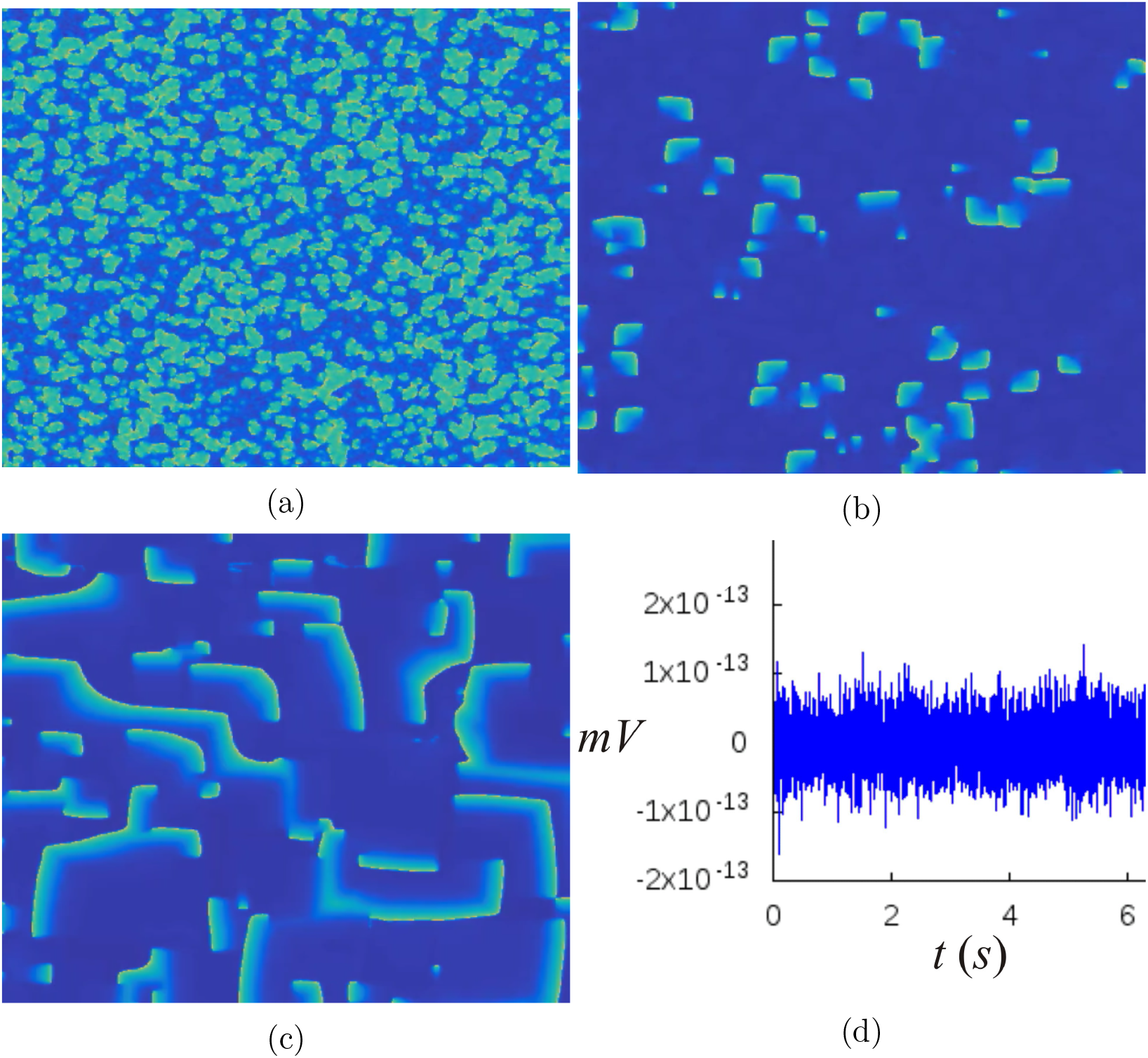
Under a different set of initial conditions than those in Figure 13, an intricate pattern of several self-generated spirals surges. (a) A large number of collisions of wavefronts after the massive blocking are apparent. (b) Some curved wavefronts survive after collisions. (c) Several spirals are generated with the curved wavefronts in (b) and persist in a steady complex state. (d) Pseudo-EG of the entire series.

A comparison of the patterns produced by the Nygren cell model and the Barkley model is shown in Figure 15. Such is the pattern produced by massive blocking in a 60× 60 net.

**Figure (15).**
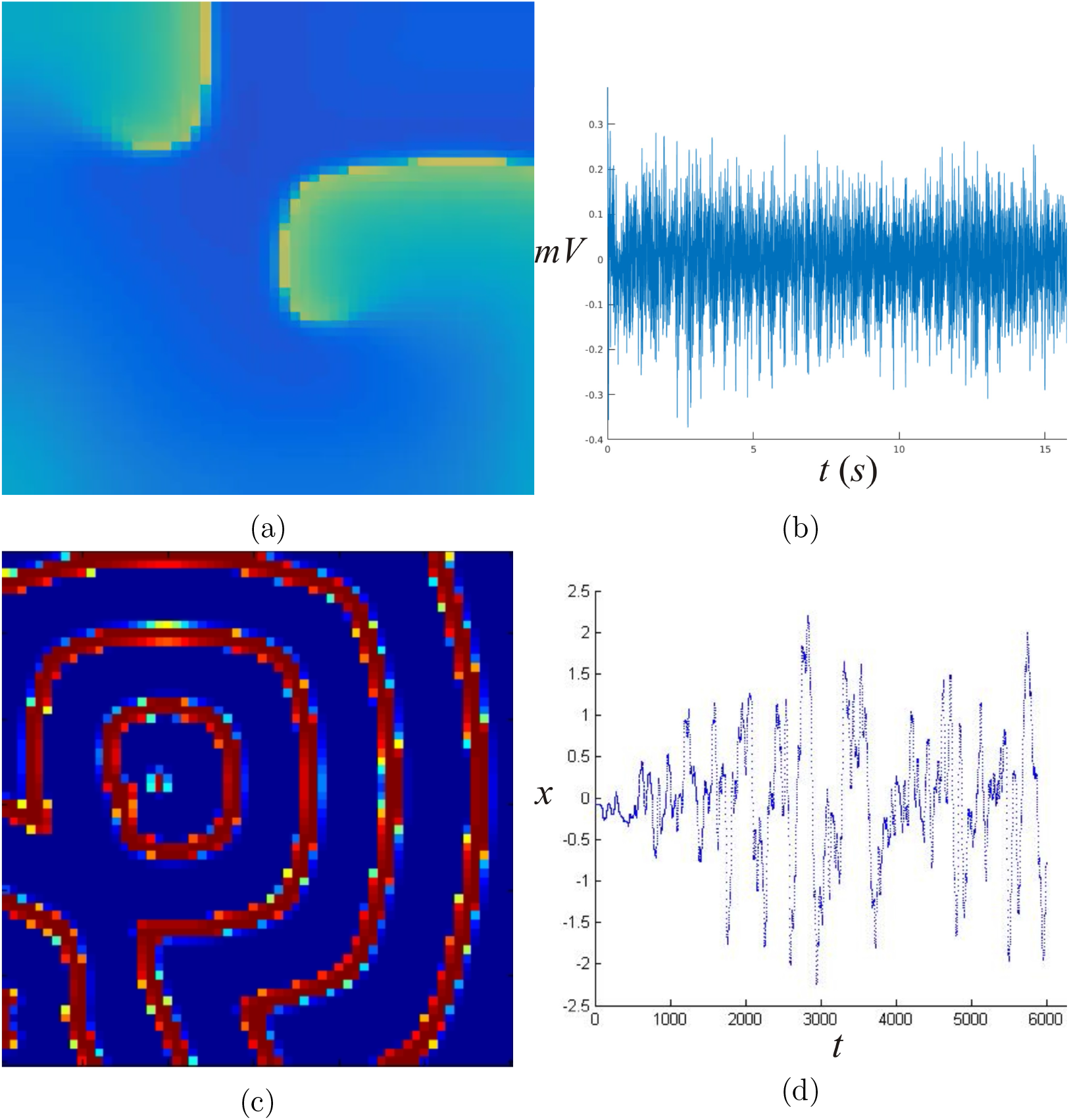
(a)Nygren model in a Fibrillation pattern. (b) Pseudo-EG of the Nygren model. (c) Barkley mode in a Fibrillation. (d) A massive blocking of individual cells with low conductivity produces an intricate pattern that, according to the pseudo-EG in the figure, can be identified with Fibrillation. Here we show systems of 60× 60 cells.

## 5. Discussion

In many articles, one and two-dimensional arrangements (for instance, in [9] and references therein, and [41]) are considered disregarding that the actual geometry of tissue is three dimensional. This reduction in modeling is a generalized attitude that can be understood under the dynamic of building models that go from the simple to the more complicated. Nevertheless, as Fenton et al., claim [10], in models of cardiac electrical activity “simulations in 3D have shown that the existence of purely three-dimensional breakup mechanisms”. So that arrhythmias are, in this sense, three-dimensional phenomena virtually. Hence, to obtain more accurate models of arrhythmias, it is necessary to work in 3D frameworks, but utilizing 2D layers makes sense for the following reasons. In modeling auricular heart tissue, it is known that rotational anisotropy of fibers of ventricular muscle can be model by superposing and rotating two-dimensional layers of cells [11]. To this aim, two ways to incorporate connectivity parameters in the cells are available: experimental histological data or stochastic or complex connectivity provided by mathematical models ad hoc. In this article, we used an aperiodic tiling model, which is possible to extend to 3D nets of cells. Anyhow, the extension to 3D of the aperiodic tiling model presented here is not straight-forward. Creating an adjacency matrix corresponding to aperiodic, fractal arrangements is a complex task and constitutes an open problem, although algorithms to produce many of such patterns are known (see, for instance, Rangel-Mondragon article [30]).

Although simplified models in one or two dimensions may reflect certain experimental data [13], it may be that such simplifications do not represent the actual behavior of big groups of cells, for instance, in the transmission of the action potential of the system formed by mixing pacemaker and atrial cells. For example, in [21] for specific mathematical models of atrial and sinus cells, the activation of the complex depends on the number of cells involved and the geometric distribution of the cells in the network. Modeling diffusive media with cells, including many observables, may not be a trivial task, since even the stability of the numerical methods involved may be challenged. Moreover, since the real geometry of the cells must be considered, real data of local topology of diffusive tissue must be incorporated, when available, to model big groups of cells. As mentioned in section 4 our representation of groups of cells in Figure 4 may not correspond to any real biological tissue, but only constitute the first approach of statistical data of a mean of the number of observed connections in the atrial heart tissue. A refinement of this data is required for the two-dimensional layers forming atrial and ventricular tissue in the heart.

On the other hand, many interesting works such as [36] study the stochastic distribution of inhomogeneities at the cellular level that can cause cardiac propagation to be stochastic. In contrast, in this article, without considering the stochastic setting, we obtained a complex propagation in the diffusive tissue, despite the simplicity of the included models, only by varying the local topology of the network. Nevertheless, variable conductance may be included in future work either employing stochastic distributions or, as was done in this article, introducing some aperiodic pattern of the distributions of the different conductances referred to in [36] of the individual cell membranes.

As a final remarck, a note about our analysis of pseudo-EG. Electrograms were statistically analyzed using the DFA technique to associate the visual behavior of the propagation of the electrical signal with a quantitative indicator of the propagation dynamics. When a scaling exponent near to 0.5 was obtained, the simulated electrical signal showed a random behavior, which corresponds to what we denominated Fibrillation (Figures 11). In the electrograms where it was possible to observe a “noisy” periodic behavior, the value of the scaling exponent *α* was near to 1, indicating long term correlations (i.e., a non-random behavior), which corresponds to the wave-like propagation observed in images and videos from the simulated experiments (Figures 8). From these results, we can conclude that images and videos of the propagation of the simulated electrical signals give us valuable information about their dynamics and factors affecting it, which is a crucial aspect to consider when analyzing structural and functional mechanisms triggering the different types of arrhythmias, such as atrial Fibrillation and Flutter.

## 6. Conclusions

In modeling cell-to-cell in this document, we found that very complex self-perpetuating diffusion patterns arise utilizing a massive blocking of cells in an excited state. This complexity emerges even in utilizing an elementary chessboard-like distribution of cells. One remarkable property of nets of diffusive cells in this document is that reentrant waves are formed in a wide variety of initial conditions contradicting the intuitive folk thinking that arrhythmia phenomena are exceptional in diffusive media, especially in considering Fibrillation.

We introduced a net with a tiling distribution in which Fibrillation, Fluttering, and a sequence of Fluttering-Fibrillation phenomena emerged. In this way, the two basic types of arrhythmia were modeled in two-dimensional tissue with a degree of complexity given by the non-periodic, fractal distribution connections in the tiling. The interesting fact is that it is possible to model a complex-like Fibrillation phenomenon by introducing a certain degree of complexity in the distribution of neighbor cells (for example, with tiles similar to those in Figure 4), instead of using any random distribution whatsoever. To the best of the knowledge of the authors of this paper, this is a novelty. Moreover, in this study, the authors found a critical value of conductivity among the cells integrating the ODE’s systems. Such critical value emerges with an adjacency matrix given by the arrangement in Figure 5. In this way, modeling Fluttering by lowing conductivity in our model of simple two-variable ODE or the state of the art ODE model by adding only specified complexity in the distribution of cells could be relevant in mathematical modeling and computational simulation.

Micro-reentry can be simulated with Barkley and Nygren models. In some in silico experiments emerged several self-perpetuating waves that collide, leading to a complex pseudo-EG, which anyhow may be identified with Fibrillation of the system. An example of this EG for such arrhythmia is shown in Figure 15(b) for the Nygren model and Figure 15(d) for Barkley model.

## Founding

This research did not receive any specific grant from funding agencies in the public, commercial, or not-for-profit sectors.

## Videos

To access all the simulation videos in this document, the reader can use the following link: http://pacifico.izt.uam.mx/aurelio/

